# scds: Computational Annotation of Doublets in Single Cell RNA Sequencing Data

**DOI:** 10.1101/564021

**Authors:** Abha S Bais, Dennis Kostka

## Abstract

**Motivation:** Single cell RNA sequencing (scRNA-seq) technologies enable the study of transcriptional heterogeneity at the resolution of individual cells and have an increasing impact on biomedical research. Specifically, high-throughput approaches that employ micro-fluidics in combination with unique molecular identifiers (UMIs) are capable of assaying many thousands of cells per experiment and are rapidly becoming commonplace. However, it is known that these methods sometimes wrongly consider two or more cells as single cells, and that a number of so-called *doublets* is present in the output of such experiments. Treating doublets as single cells in downstream analyses can severely bias a study’s conclusions, and therefore computational strategies for the identification of doublets are needed. Here we present single cell doublet scoring (`scds`), a software tool for the *in silico* identification of doublets in scRNA-seq data.

**Results:** With `scds`, we propose two new and complementary approaches for doublet identification: Co-expression based doublet scoring (`cxds`) and binary classification based doublet scoring (`bcds`). The co-expression based approach, `cxds`, utilizes binarized (absence/presence) gene expression data and employs a binomial model for the co-expression of pairs of genes and yields interpretable doublet annotations. `bcds`, on the other hand, uses a binary classification approach to discriminate artificial doublets from the original data. We apply our methods and existing doublet identification approaches to four data sets with experimental doublet annotations and find that our methods perform at least as well as the state of the art, but at comparably little computational cost. We also find appreciable differences between methods and across data sets, that no approach dominates all others, and we believe there is room for improvement in computational doublet identification as more data with experimental annotations becomes available. In the meanwhile, `scds` presents a scalable, competitive approach that allows for doublet annotations in thousands of cells in a matter of seconds.

**Availability and Implementation:** `scds` is implemented as an R package and freely available at https://github.com/kostkalab/scds.

## Introduction

Single cell RNA sequencing (scRNA-seq) technologies allow characterization of transcriptomes of individual cells, and aid our understanding of tissue and cell-type heterogeneity. New insights, for instance in the context of development and/or disease ((Li *et al.*, 2017; Segerstolpe *et al.*, 2016); for a review see (Potter, 2018)), have made them increasingly relevant across a diverse range of biomedical research fields. Specifically, micro-fluidics based approaches utilize unique molecular identifiers (UMIs) and enable the study of many thousands of cells simultaneously. The availability of user-friendly solutions (like the 10X Chromium platform) has rendered this flavor of UMI-based scRNA-seq the assay of choice in numerous studies. However, use of scRNA-seq data is not without challenges, and careful data processing, quality control and analysis is essential (reviewed for instance in (Stegle *et al.*, 2015; Vallejos *et al.*, 2017; AlJanahi *et al.*, 2018; Kiselev *et al.*, 2019). We focus on one key step of quality control that is the identification of so-called *“doublets/multiplets”*. Doublets (or multiplets) arise in scRNA-seq data when two (or more) cells are mistakenly considered as a single cell, due for instance to being captured and processed in the same droplet on a micro-fluidics device. This type of error has the potential to severely confound interpretation of study results, especially in the context of cellular heterogeneity and identity, where they may appear as spurious novel cell types. However, despite rapid advances in the field, to our knowledge relatively few approaches exist that address the issue of doublet detection in scRNA-seq data. In the following, we provide a brief overview of existing experimental and computational approaches for doublet identification.

### Experimental methods for doublet detection

For some approaches doublet detection can be performed as a quality control step to ensure that only single cells are picked at capture sites (e.g. (Proserpio *et al.*, 2016; Segerstolpe *et al.*, 2016)). Alternatively, barcodes have been used together with mixtures of cells from different species to get estimates of doublet rates (e.g., (Klein *et al.*, 2015; Alles *et al.*, 2017)). In their work, (Kang *et al.*, 2018) present a multiplexing strategy that exploits genetic variation to detect doublets among mixtures of cells from different individuals. In another approach, (Stoeckius *et al.*, 2018) use oligonucleotide-tagged antibodies against cell surface proteins to uniquely label cells in a robust multiplexing strategy that allows for doublet detection. In a similar vein, (Gehring *et al.*, 2018) use chemical labeling for tagging cells from individual samples. Recently, (McGinnis *et al.*, 2018) proposed a technique called MULTI-seq, which uses lipid-modified oligonucleotides to barcode individual cells. Thus, development of experimental approaches that improve doublet detection is a field of active research. However, experimental approaches typically face the limitation that they require specific technologies or experimental designs, which are often not readily available to researchers (for an overview of limitations of some of these approaches see (Wolock *et al.*, 2018)). Therefore, it is at the stage of computational data analysis where approaches are needed to identify doublets.

### Computational methods for doublet detection

There are few computational approaches that explicitly address the problem of distinguishing doublets from single cells using scRNA-seq expression data alone. Often, researchers rely on curated marker genes and expert knowledge to identify cells co-expressing markers of distinct cell types as putative doublets (e.g., (Wang *et al.*, 2016; Ibarra-Soria *et al.*, 2018; Rosenberg *et al.*, 2018)). Based on the assumption that doublets would have higher total RNA content, another approach is to use a measure for overall expression signal (total counts, for example) as a means for classifying cells as doublets (Bach *et al.*, 2017; Ziegenhain *et al.*, 2017; Krentz *et* al., 2018). However, given that marker gene information and expert knowledge is not always available (and not always objective), and that doublets may not necessarily have high total counts, in the last year a number of computational doublet detection/annotation methods have been proposed that do not rely on markers or total counts alone (Lun *et al.*, 2016; Shor and Gayoso, 2019; Wolock *et al.*, 2018; McGinnis *et al.*, 2018; DePasquale *et al.*, 2018), see Table 4. In the following we briefly summarize each of them:

`scrublet`: In their approach `scrublet`, (Wolock *et al.*, 2018) simulate artificial doublets from the original data coordinates of the normalized and filtered data in a reduced-dimensional representation obtained by principal component analysis (PCA). A *doublet score* is then created by considering the fraction of artificial doublets in the neighborhood of each barcode using k-nearest-neighbor (kNN) graph based on Euclidean distances. To determine the fraction of doublets in an experiment, a doublet score threshold is set visually by comparing the distributions of the doublet scores of original barcodes and artificial doublets. `scrublet` is available as a python module.

`dblFinder`: In a similar vein, `DoubletFinder` (McGinnis *et al.*, 2018) also uses artificial doublets, and the fraction of artificial doublets in the neighborhood of each barcode, to calculate a metric (“pANN”), akin to the doublet score discussed above. Artificial doublets are created by averaging raw counts of randomly paired barcodes, then the data is normalized, PCA is performed, and pANN scores are computed. The authors provide a heuristic to automatically choose parameters (like the number of neighbors considered), and finally thresholding pANN based on the expected doublet rate (or base on an adjusted rate that accounts for homotypic doublets (doublets formed by cells of the same type)) yields final doublet annotations. `dblFinder` is available as an R package.

`dblCells`: In the vignette of their R package `simpleSingleCell` Lun et al. 2016 discuss two approaches, `doubletClusters` and `doubletCells`, implemented as part of the R package `scran` (Lun 2016). The first prescribes an approach to identify clusters of cells that have intermediate expression profiles to “parent” clusters based on differentially expressed genes, library size and number of cells in a cluster (Bach et al. 2017). Of relevance to us is the second approach `doubletCells`, whereby thousands of artificial doublets are generated by combining randomly chosen pairs of barcodes and projecting them into a reduced-dimensional space. A doublet score is formalized by assessing neighborhoods of simulated doublets and original barcodes.

`dblDetection`: This approach (JonathanShor/DoubletDetection) also relies on artificially generated doublets, but, in contrast to previous methods, performs cell clustering on the augmented data set. Briefly, augmented data with artificial doublets is generated from one of two possible sampling schemes, projected into a lower-dimensional representation using PCA, and then clustering is performed with phenograph (Levine *et al.*, 2015; JonathanShor/PhenoGraph). Next, hypergeometric p-values are assigned to clusters and their cells based on the number of artificial doublets they contain. This procedure (including artificial doublet generation) is performed multiple times, and then doublet calls and scores are derived from annotated p-values across runs/iterations. `dblDetection` is available as a python module.

`dblDecon`: Making use of an initial user-provided clustering, the method of (DePasquale *et al.*, 2018), DoubletDecon, relies on deconvolution as implemented in the R package DeconRNASeq (Gong and Szustakowski, 2013) to identify doublets. First, distinct reference profiles are constructed from the initial clustering, and then artificial doublets are generated and their deconvolution profiles are computed. Next, barcodes with deconvolution profiles closest (by Pearson correlation) to those of a synthetic doublet are initially predicted to be a doublet. Finally, to reduce penalizing cells with gene expression profiles possibly corresponding to transitional cell states, the authors implement a “Rescue” step whereby predicted doublets with unique gene expression patterns are re-labeled as single cells. `dblDecon` is available as an R package.

We note that most of these approaches are recent, based on similar strategies, and to our knowledge have not been assessed together across multiple data sets in a systematic way. In the following we present two new and complementary methods for computational doublet annotation: Co-expression based doublet scoring (`cxds`) and binary classification based doublet scoring (`bcds`). We show that they can accurately annotate doublets, and we perform a comparison of these approaches and the methods discussed above on four publicly available data sets with experimental doublet annotations (see **Table 1**). We show that our methods perform well compared with existing approaches (at comparably little computational cost), and we demonstrate heterogeneity in results and performance of computational doublet annotation between different methods and across different data sets.

**Table 1:** Data sets with experimental doublet annotation. The “# Genes” column shows the median across cells of the number of expressed genes

| Data set | Cells | Sparsity | # Genes | Annotation type | Reference |
| --- | --- | --- | --- | --- | --- |
| hgmm | 12,820 | 79% | 3,068.5 | cross species cell line mixture | (10X_Genomics) |
| ch_pbmc | 15,583 | 98% | 321 | cell hashing | (Stoeckius *et al.*, 2018) |
| ch_cell-lines | 8,191 | 92% | 2,086 | cell hashing | (Stoeckius *et al.*, 2018) |
| demuxlet | 14,619 | 97% | 520 | genetic variation | (Kang *et al.*, 2018) |

## Materials and methods

### Co-expression based doublet scoring

Co-expression based doublet scoring (`cxds`) is motivated by the assumption that heterotypic doublets (i.e., doublets comprised of cells from different cell types), co-express “marker” genes that are not usually active in the same cell. In contrast to approaches that leverage expert knowledge and assess expression patterns of curated sets of marker genes manually, `cxds` uses the available data to first assess gene pairs and then derive an overall doublet score for each barcode^1^, based on gene-gene co-expression.

Specifically, let *X* ∈ ℝ^*m*×*n*^ be a genes x cells count matrix for *m* genes and *n* cells, and *B* its thresholded binarized version, where *B*_*ij*_ denotes whether gene *i* is expressed in cell *j* (absence/presence). The row means of *B*, 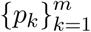, are the fraction of cells expressing each gene, and the symmetric matrix *BB*^*T*^ contains for each gene pair the number of cells co-expressing the two genes. If we denote the matrix where the entries in *B* have been flipped by *<u>B</u>* (*<u>B</u>*_*ij*_ = 1 − *B*_*ij*_), then we can write the number of cells that express exactly one of two genes as (*<u>B</u>B*^*T*^ + *<u>B</u>B*^*T*^) and, assuming independence between genes, arrive at the following binomial model:

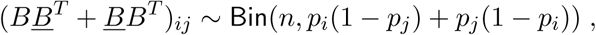

where (*ij*) now denotes a pair of genes. Let a “score matrix” *S* ∈ ℝ^*m×m*^ hold negative (upper tail) log p-values under the above model. Scores for gene pairs that co-express across cells **less often** than expected (given their marginal frequencies) are high, while scores for pairs that co-express normally (or more often than expected) are low. We now use *S* to derive cell-specific doublet scores by summing, for each cell, negative log p-values of co-expressed gene pairs, so that we get for cell *i* a doublet score `cxds` via:

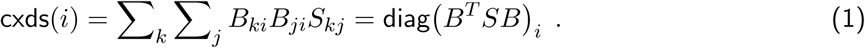

We then rank cells in the order of decreasing scores, with high scores denoting doublet cells/barcodes. We note that *B*, for UMI data, is typically sparse (often more than 95% zeros), so that the matrix products *BB*^*T*^ and *B*^*T*^ *SB* above are not prohibitive, even for tens of thousands of cells. On the contrary, our run times are comparable with the fastest current approaches (see **Results** section).

As mentioned above, a motivation for this score is a (simplified) concept of marker genes that are expressed in specific cell types only. Gene pairs containing marker genes for the same cell type will receive low scores (they are co-expressed more often than expected), while gene pairs with marker genes for different cell types would receive high scores (they are co-expressed less often than expected, because they do not co-express in non-doublet cells). In our cell-specific scores 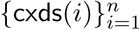 we then aggregate information across gene pairs.

#### Gene pair scoring

Because the doublet score cxds(·) in Equation (1) directly sums up contributions of individual gene pairs, we can rank pairs based on their cumulative impact on doublet prediction in the data set at hand, weighted by the doublet score for each cell. For the “importance” of a pair formed by genes *k* and *j* we define

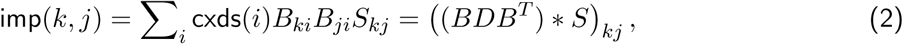

where *D* is a diagonal matrix containing doublet scores and ∗ denotes the element-wise product of matrices. This approach prioritizes gene pairs that substantially contribute to the annotation of cells with high doublet scores, and it can be used to study the pairs of genes that most drive doublet prediction. Further on, to prioritize gene pairs that drive doublet predictions in a particular cell we can omit the sum in Equation (2); or, to focus on a group of cells (forming a cluster, for instance), we can restrict the sum to group members.

#### Implementation

We implemented `cxds` using the R programming language (R Core Team, 2018), and in practice add two heuristics: Given a count matrix *X* of an scRNA-seq experiment, we first binarize expression based on a threshold `binThresh`, such that *B* contains genes with more than `binThresh` counts. In all our studies here we set `binThresh` to zero, but other values can be reasonable. Next, we focus on highly variable genes by ranking genes with respect to their Binomial variance (i.e., *np*_*j*_(1 *-p*_*j*_) for gene *j*) then keeping only the `ntop` most variable ones. We choose `ntop=500` as default.

### Binary classification based doublet scoring

Binary classification based doublet scoring (`bcds`) employs artificial doublets, similar to other strategies (see the **Introduction** for an overview). However, it does not rely on dimension reduction or nearest neighbor approaches to calculate a doublet score. Briefly, given a genes-by-cells matrix of expression counts we create artificial doublets by adding random pairs of columns. We then log-transform, normalize, and select variable genes before using a binary classification algorithm to discriminate artificial doublets from original input data. Finally, for each input barcode we then take the estimated probability of belonging to the artificial doublet class as the doublet score we annotate.

#### Implementation

We implemented `bcds` using the R programming language (R Core Team, 2018), with the following specifics. We simulate artificial doublets by randomly selecting pairs of cells and adding their counts, followed by mean-normalization of log-counts of all cells (artificial doublets and input cells) and thereby generate an augmented dataset containing input data and simulated doublets. We then train gradient boosted decision trees (Chen and Guestrin, 2016) using the `xgboost` R package (Chen *et al.*, 2019) with default parameters for artificial doublet classification. We employ two heuristics for establishing the number of training rounds: (i) We use 5-fold cross-validation approach in combination with the “one-standard-error-rule” (Hastie *et al.*, 2001) to determine the number of rounds to train on the complete data set. (ii) We set the number of training rounds to seven. In both cases we stop training in case the misclassification error does not decrease for two consecutive rounds. All results reported in this manuscript use heuristic (i), except for **Table 3**, where we report running times; there we also report heuristic (ii), termed `bcds_7` (supplemental **Table 7** compares the performance of the two heuristics across data sets). We report the class probability for the artificial doublet class given by the model trained on the complete data set as doublet scores. Also, like with `cxds`, we select `ntop` variable genes before simulating doublets and training the classifier. Here, we log-transform and mean-normalize count values before calculating the variance of each gene. The `ntop` most variable genes are then included for further analysis, and we choose `ntop=500` for all results reported.

**Table 2:** Performance of doublet annotation methods, averaged across data sets. Gray bullets mark baseline methods, blue bullets mark current methods for doublet annotation, and red bullets mark proposed methods.

|  | AUROC | pAUC90 | pAUC95 | pAUC97.5 | AUPRC |
| --- | --- | --- | --- | --- | --- |
| • dblCells | 0.75 | 0.69 | 0.66 | 0.63 | 0.47 |
| • libsize | 0.76 | 0.60 | 0.56 | 0.53 | 0.30 |
| • features | 0.78 | 0.62 | 0.57 | 0.54 | 0.33 |
| • cxds | 0.83 | 0.74 | 0.71 | 0.68 | 0.56 |
| • scrublet | 0.83 | 0.76 | 0.72 | 0.69 | 0.59 |
| • bcds | 0.84 | 0.76 | 0.70 | 0.64 | 0.57 |
| • dblDetection | 0.84 | 0.80 | 0.75 | 0.71 | 0.65 |
| • hybrid | 0.85 | 0.77 | 0.72 | 0.67 | 0.62 |
| • dblFinder | 0.86 | 0.80 | 0.76 | 0.72 | 0.68 |

**Table 3:** Running time for doublet detection methods in seconds. Missing values indicate running times longer than 300 seconds (5 minutes).

|  | 1,000 | 2,000 | 4,000 | 8,000 | 12,000 |
| --- | --- | --- | --- | --- | --- |
| bcds_7 | 5 | 2 | 4 | 11 | 9 |
| cxds | 1 | 1 | 3 | 9 | 21 |
| scrublet | 5 | 6 | 10 | 16 | 21 |
| bcds | 12 | 9 | 14 | 74 | 41 |
| dblCells | 52 | 29 | 170 | 276 | × |
| dblFinder | 93 | 136 | 269 | × | × |
| dblDetection | × | × | × | × | × |

### Hybrid doublet scoring

We also combine both approaches, `cxds` and `bcds`, into a version generating annotations as follows. After running each method we simply normalize the scores to fall between zero and one (by subtracting the minimum and subsequently dividing by the maximum) before adding them. We denote these annotation scores as `hybrid`.

### Data description, retrieval and processing

We evaluated our aproach and compared performance with other methods on four publicly available data sets with experimentally annotated doublets. **Table 1** lists the data sets, in the following we describe how we retrieved and processed each of them:

`hg-mm`: This data set contains a 1:1 mixture of freshly frozen human HEK293T cells and mouse NIH3T3 cells. We downloaded data from the 10X genomics website (10X Genomics) and processed as follows: Barcodes were filtered to include those with experimental doublet annotations. For genes, human-mouse 1:1 orthologs were identified using the Ensemble database (v95, (Zerbino *et al.*, 2018)) with the `getLDS` function provided by the `biomaRt` R software package (Durinck *et al.*, 2009), and corresponding counts were added. Removing features with no counts resulted in gene expression data of 14,437 orthologs across 12,820 barcodes.

`ch_pbmc`: This data set contains peripheral blood mononuclear cells (PBMCs) from eight donors, with cells from each donor uniquely labeled using the cell hashing approach of (Stoeckius *et al.*, 2018). Data files were downloaded from Dropbox (url) and processed according to the vignette of the `Seurat` R package (Butler *et al.*, 2018) entitled “Demultiplexing with hashtag oligos (HTOs)” (url). This resulted in a gene expression matrix of 21,606 genes across 15,583 barcodes.

`ch_cell-lines`: This data set contains a mixture of four human cell lines, HEK, K562, KG1 and THP1. Each cell line was labeled using the cell hashing approach of (Stoeckius *et al.*, 2018). Data files were downloaded from the same location as for `ch_pbmc` and processed according to the same vignette, resulting in a gene expression matrix with 25,241 genes across 8,191 barcodes.

`demuxlet`: This data set contains a uniform mixture of PBMCs from eight lupus patients, and doublets have been annotated based on genetic information using `demuxlet` (Kang *et al.*, 2018). Data files for gene expression counts were downloaded from GEO (GSM2560248) and doublet annotations were trieved from the `demuxlet` github repository (url). This resulted in data comprising of expression counts for 17,662 genes across 14,619 barcodes.

We note that for all gene counts above, and for the sparsity calculations in **Table 1**, we included genes expressed with at least one count in one barcode.

### Annotation of doublets with existing methods

We annotated doublets with five existing tools (**Table 4)**, and in the following we describe how we applied each of them:

**Table 4:**
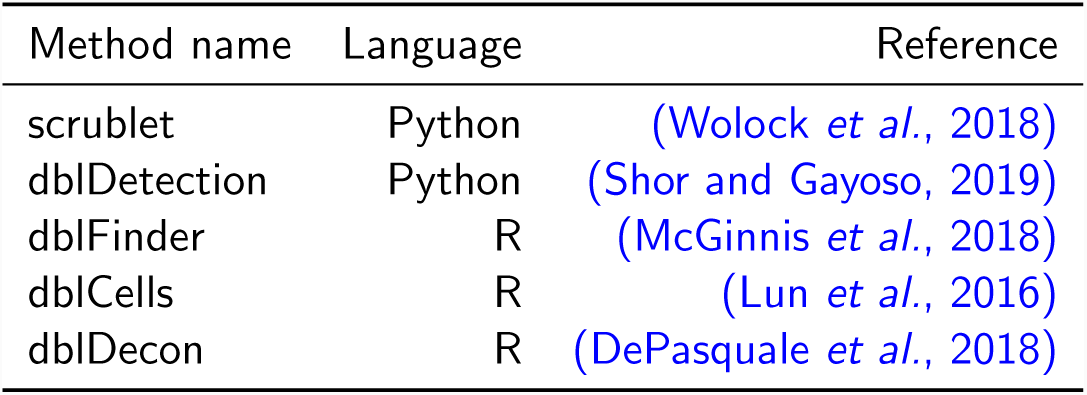
Computational methods for doublet annotation summarized in Introduction.

| Method name | Language | Reference |
| --- | --- | --- |
| scrublet | Python | (Wolock *et al.*, 2018) |
| dblDetection | Python | (Shor and Gayoso, 2019) |
| dblFinder | R | (McGinnis *et al.*, 2018) |
| dblCells | R | (Lun *et al.*, 2016) |
| dblDecon | R | (DePasquale *et al.*, 2018) |

`dblCells`: Data was processed per the vignette of the R package `simpleSingleCell`(Lun et al. 2016). Briefly, raw counts were normalized using size factors computed using `scran` (Lun 2016) with the `igraph` clustering method and a `min.mean` value of 0.1. Technical noise was removed using the `denoisePCA` function of `scran` with approximate singular value decomposition performed (`approximate = TRUE`). Finally, doublet scores were retrieved using the `doubletCells` function run with default options except again with `approximate = TRUE` to allow fast approximate PCA.

`dblDecon`: Raw counts were fully processed using Seurat (Butler *et al.*, 2018) (i.e. normalization, scaling with nUMI regressed out, finding variable genes, dimension reduction (with PCA) and clustering were performed). Additionally, marker genes were calculated with default settings using the `FindAllMarkers` function and top 50 markers used. The `Main Doublet Decon` function was run with input files created using the `Seurat Pre Process` function and default settings except for `species` which was set to `hsa`, and using centroids as references for deconvolution (`centroids = TRUE`).

`dblDetection`: The python module (JonathanShor/DoubletDetection) was used in the R programming language using the `reticulate` package (Allaire *et al.*, 2018), and run with default parameters on the count data. For each cell, negative log p-values were averaged across runs/iterations to derive an aggregate doublet score per cell.

`dblFinder`: Fully processed Seurat (Butler *et al.*, 2018) object was created, where normalization, scaling (with nUMI regressed out), finding variable genes (with arguments as per their github example code), dimension reduction (PCA and TSNE) and clustering were performed with `dims.use = 10` and all other Seurat settings set to default. For `dblFinder`, the value for `pk` was selected following the best practices outlined on their github page (McGinnis, 2018), as the one at which the mean-variance normalized bimodality coefficient (BCmvn) is maximized. The function `DoubletFinder` was run with the expected doublet rate of 7.5% assuming Poisson statistics, as per the example code on github (McGinnis, 2018).

`scrublet`: The python module (AllonKleinLab/scrublet) was used in the R programming language using the `reticulate` package (Allaire *et al.*, 2018), and run with default parameters on raw count data. Doublet scores were used as reported by the software.

### Data visualization and calculation of performance metrics

Low dimensional representation for visualization of data in our figures were calculated as follows: For each data set, log counts were calculated and random projection PCA was performed on the 500 most variable genes using the `rsvd` R package (Erichson *et al.*, 2016); finally the first ten principal components were projected into two dimensions for visualization using the `Rtsne` package (Krijthe, 2015) with default parameters.

For performance evaluation we calculated the area under the ROC curve using the `pROC` R package (Robin *et al.*, 2011), including partial areas under the ROC curve (pAUC) at 90%, 95% and 97.5% specificity. For the partial areas, the option `partial.auc.correct` was set to `TRUE`, such that the maximal pAUC is one and a pAUC of 0.5 is non-discriminant. Areas under the precision-recall curves (AUPRCs) were calculated using the `PRROC` package (Keilwagen *et al.*, 2014) and we report the smoothed area under the curve according to (Davis and Goadrich, 2006) by selecting the appropriate option. We used all cells present in each data set (see above) to calculate performance metrics. We note that `dblDetection` would occasionally not score a small subset of cells (between zero and eleven), which we then excluded for this method’s metrics.

Running times were calculated in R using the `reticulate` package (Allaire *et al.*, 2018) for python modules, and the median (middle) value of three timings is reported. For timings, methods were run on the same sub-samples of cells of the `demu` data, and two cores of an Intel(R) Xeon(R) E5-2667 v4 CPU cpu were made available for computing.

## Results

We report two computational methods for *in silico* doublet prediction: co-expression based doublet scoring (`cxds`) and binary classification based doublet scoring (`bcds`). Co-expression based doublet scoring identifies doublets from thresholded expression data essentially using a Binomial model (see **Methods** section), based on the reasoning that marker genes for different cell types do not co-express in (non-doublet) barcodes. Pairs are scored exhaustively, and no prior knowledge about marker genes in a specific context is needed. Further on, doublet annotations for cells are interpretable in the sense that they are based on the co-expression of pairs of genes, and `cxds` allows users to view gene pairs ordered with respect to their contribution to doublet predictions across a data set (see **Methods** section). **Figure 1** shows the five top gene pairs driving doublet prediction for `cxds` across the four data sets (**Table 1**), illustrating how co-expression of genes in each pair identifies doublet cells.

Binary classification based doublet scoring, on the other hand, combines generation of artificial doublets from existing data with binary classification. Barcodes in the original data that are difficult to discriminate from artificial doublets receive high doublet scores using `bcds`. We use gradient boosted decision trees (Chen and Guestrin, 2016) to classify (see **Methods** section), but in principle the approach is generic and other classification algorithms could be explored. In the following, we apply our methods to four data sets with experimental doublet annotations (**Table 1)**, provide evidence that combining the `cxds` and `bcds` into a “`hybrid`” score (see **Methods** section) improves performance, and compare our approaches and other computational methods for doublet annotation (**Table 4).**

### Doublet scoring with scds accurately recapitulates experimental doublet annotations

We performed computational doublet annotation on four scRNA-seq data sets (**Tables 1 and 2**) using several current methods (**Table 4**), together with library size (`libsize`) and number of expressed genes (termed `features`; together both are referred to as “baseline” methods from here onwards). Results are summarized in **Table 5**, where columns are performance metrics (the area under the receiver operating characteristic curve (ROC curve), the area under the precision-recall curve (PR curve), and partial areas under the ROC curve focusing on 90%, 95% and 97.5% specificity), while rows correspond to computational doublet annotation approaches. For each data set, rows are sorted with respect to their performance in terms of the area under the ROC curve (AUROC), with ties being broken by the performance in terms of the area under the PR curve (AUPRC). Baseline methods are marked with gray bullets, current methods with blue bullets, and our proposed approaches with red bullets. We find that all the methods we propose (`cxds`, `bcds` and `hybrid`) perform well across data sets, consistently outperforming baseline approaches and at least one ranking in the top three best performing methods. The one exception is the `ch_pbmc` data set, where annotating doublets based on the number of features achieves an area under the ROC curve of 79%. Our weakest-performing approach on this data, `cxds`, performs slightly worse (78%), but does much better in terms of area under the PR curve (AUPRC of 54% vs 45%, respectively). We also note that two other computational doublet annotation methods, `dblCells` and `scrublet`, perform worse than the number of features in terms of AUROC on this dataset. On average, our `hybrid` method does best of the three methods we propose, significantly outperforming baseline approaches on all four data sets.

### Co-expression based doublet scoring highlights informative gene pairs

One of the features of `cxds` is its ability to provide gene pairs that drive doublet annotations of cells in a specific data set (see **Methods**). As an illustration, **Figure 1** shows the top five gene pairs (that span a unique set of ten genes so that no gene appears twice) driving `cxds` doublet annotation in each of the data sets we analyzed. For each data set (panels A - D), the first row shows a two-dimensional representation of all cells (left), the subset of experimentally annotated doublets (middle) and the subset of doublets predicted by `cxds` (right). The next five rows depict gene pairs: Binarized expression (presence/absence) of one gene alone on the left, of the second gene in the pair in the middle, and co-expression of both genes in the same cell on the right (also absence/presence). We see that `cxds` finds genes with complementary expression patterns that mark coherent groups of cells, and how coexpression of these genes contributes to doublet predictions. We note that while no clustering has been performed, genes included in high-scoring pairs by `cxds` often look like they mark different cell types, or combinations thereof, that may be present in the data.

**Figure 1:**
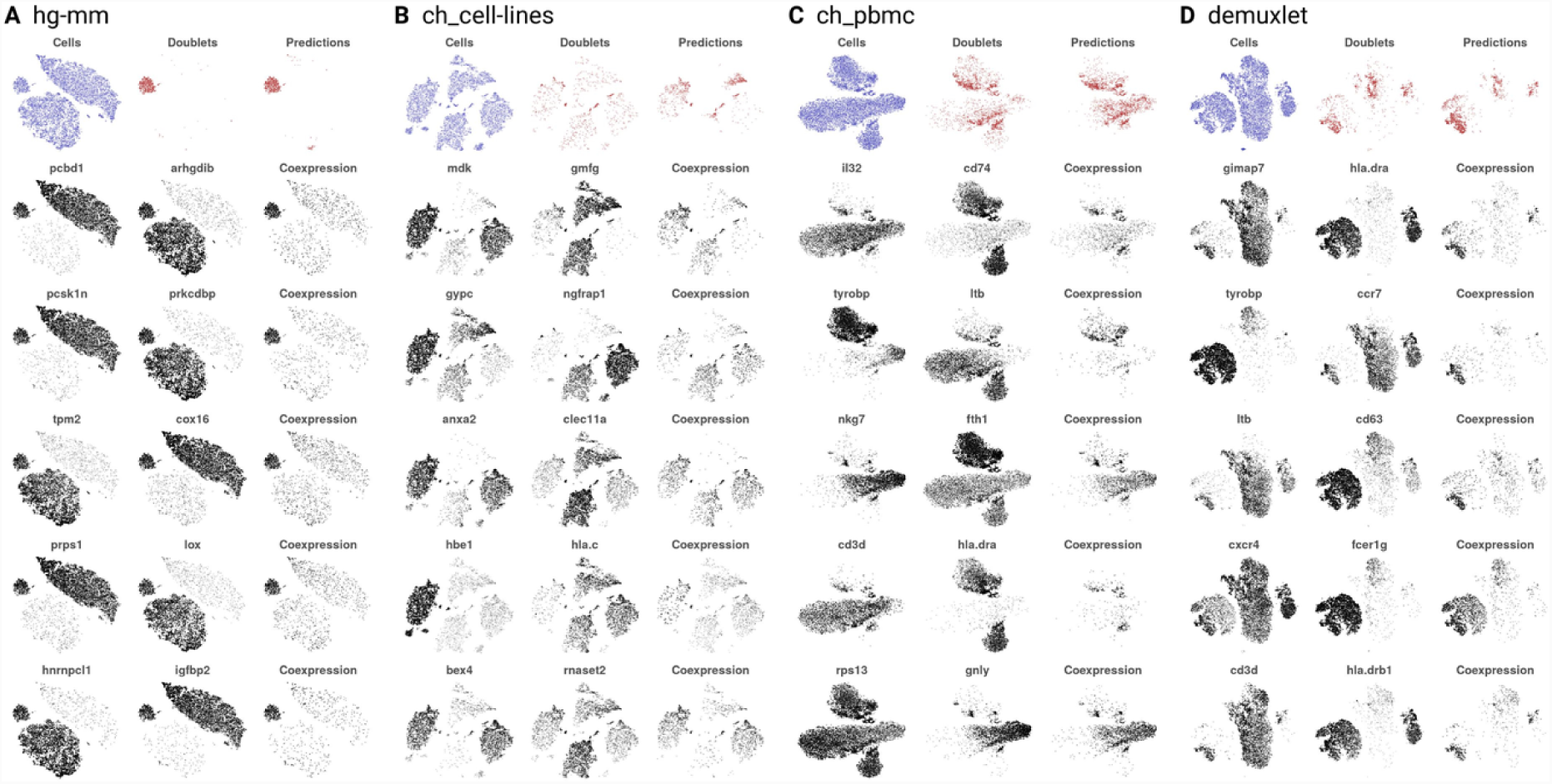
Gene pairs driving doublet prediction in cxds. For four data sets (panels A-D) the first row shows all cells in blue (left), the annotated doublets in red (center) and cxds-predicted doublets (right), also in red. The following five rows depicts the five gene pairs that contribute most to the cxds classifier (see Methods). For each pair (i.e., for each row), the left plot depicts the expression of one gene (darker = higher expression), the middle plot the expression of the other gene, while the right plot the average expression in cells that co-express both genes. We see that each gene in a pair is expressed in distinct groups of cells, and that their co-expression highlights annotated and predicted doublets.

### Comparison of computational doublet scoring methods

We compared computational doublet annotation methods across four data sets, and results are shown in **Table 5**, and average performance across data sets is summarized in **Table 2**. We find that computational doublet prediction performs best on the `hg-mm` data set, followed by `demuxlet` and `ch_pbmc`, while it is most challenging for the `ch_cell-lines` data. Within each data set there is appreciable spread of performance between the different methods, with most methods consistently outperforming baseline approaches. From **Table 2** we see that, on average, `dblFinder` performs best, followed by our `hybrid` approach and `dblDetection`. However, this order varies between data sets; for example on the `ch_cell-lines` data set, `bcds` performs better than `hybrid` **(Table** 5). On the `ch_pbmc` data set, the baseline classifiers do better than on other data sets, outperforming `cxds`, `scrublet` and `dblCells`. In general, library size and number of features identify doublets reasonably well (AUCs ≥ 78%, with the exception of the `ch_cell-lines` data set), which motivated us to further explore the effect of library size on the performance of computational doublet annotation.

#### Doublet annotation performance stratified by library size

For each data set, we divided cells into equal-sized bins according to library size, so that the first bin contains cells with library sizes between the 0% and 10% quantile, the second bin cells between the 10% and 20% quantile, and so on. We then assessed annotation performance for all computational methods in each bin for each data set separately. Results are summarized in **Figure 2**. Major columns correspond to data sets, and for each data set two panels are shown (rows). The first row depicts performance in terms of the area under the ROC curve (AUROC), the second row in terms of the area under the PR curve (AUPRC). For each performance comparison, columns correspond to library size bins, and rows to annotation methods.

**Figure 2:**
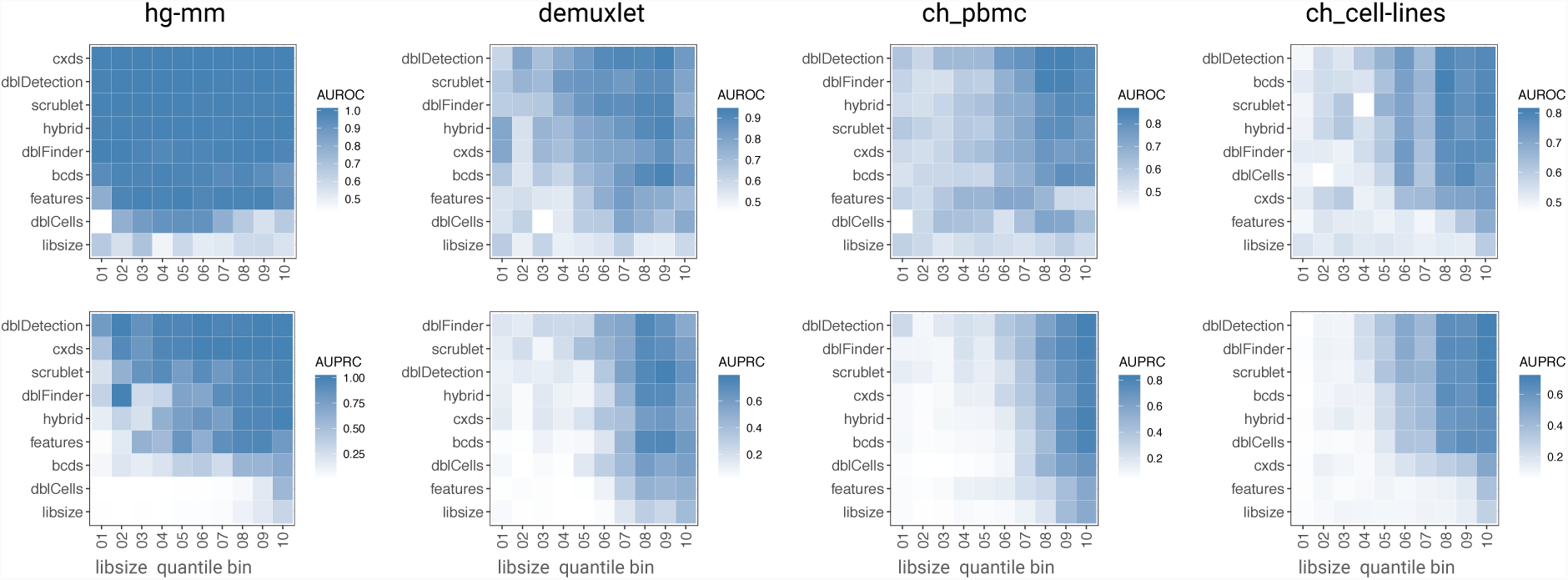
Performance of methods, stratified by library size. For each data set (columns), the first row of panels shows performance in terms of the area under the ROC curve (AUROC), while the second row shows performance under the precision-recall curve (AUPRC). For each panel the rows correspond to methods, and the columns to groups of cells in the same stratum of library sizes. The left-most column focuses on the 10% of cells with the lowest library size, the next column on the cells between the 10% and the 20% quantile, etc. In each panel methods are ranked by their average performance across quantile bins.

We see that all approaches (baseline methods included) perform best on cells with high library size (quantile bins 5 and up), and that this trend is more pronounced for performance in terms of AUPRC, compared with AUROC. We also find that this trend applies broadly, with notable exceptions being: The `hg-mm` data set, where most methods perform well in terms of AUROC across bins, and `dblDetection` and `cxds` both also perform consistently across a wide range of bins in terms of AUPRC. The second exception is the `demuxlet` data set, where `hybrid` and `cxds` perform unusually well in terms of AUROC for cells with small library sizes (first quantile bin).

#### Comparison of doublet annotations between methods

We also assessed similarities and differences between doublet predictions of each method. To do so, we determined the fraction of barcodes experimentally annotated as doublets and then compared the same number of doublet predictions for each method. Results are summarized in **Figure 3**, where we looked at overlapping and non-overlapping doublet annotations in the form of upset plots (Conway *et al.*, 2017) for each data set. Vertical bars indicate the number of cells in each intersection class described in the lower portion of the plot. Set sizes are the number of experimentally annotated doublets (i.e., they are identical across methods). Gray bars correspond to intersections containing barcodes not annotated as doublet (i.e., false positives (FP)), whereas black bars correspond to barcodes annotated as doublets (i.e., true positives (TP)). The twenty largest intersection sets are shown for each data set.

**Figure 3:**
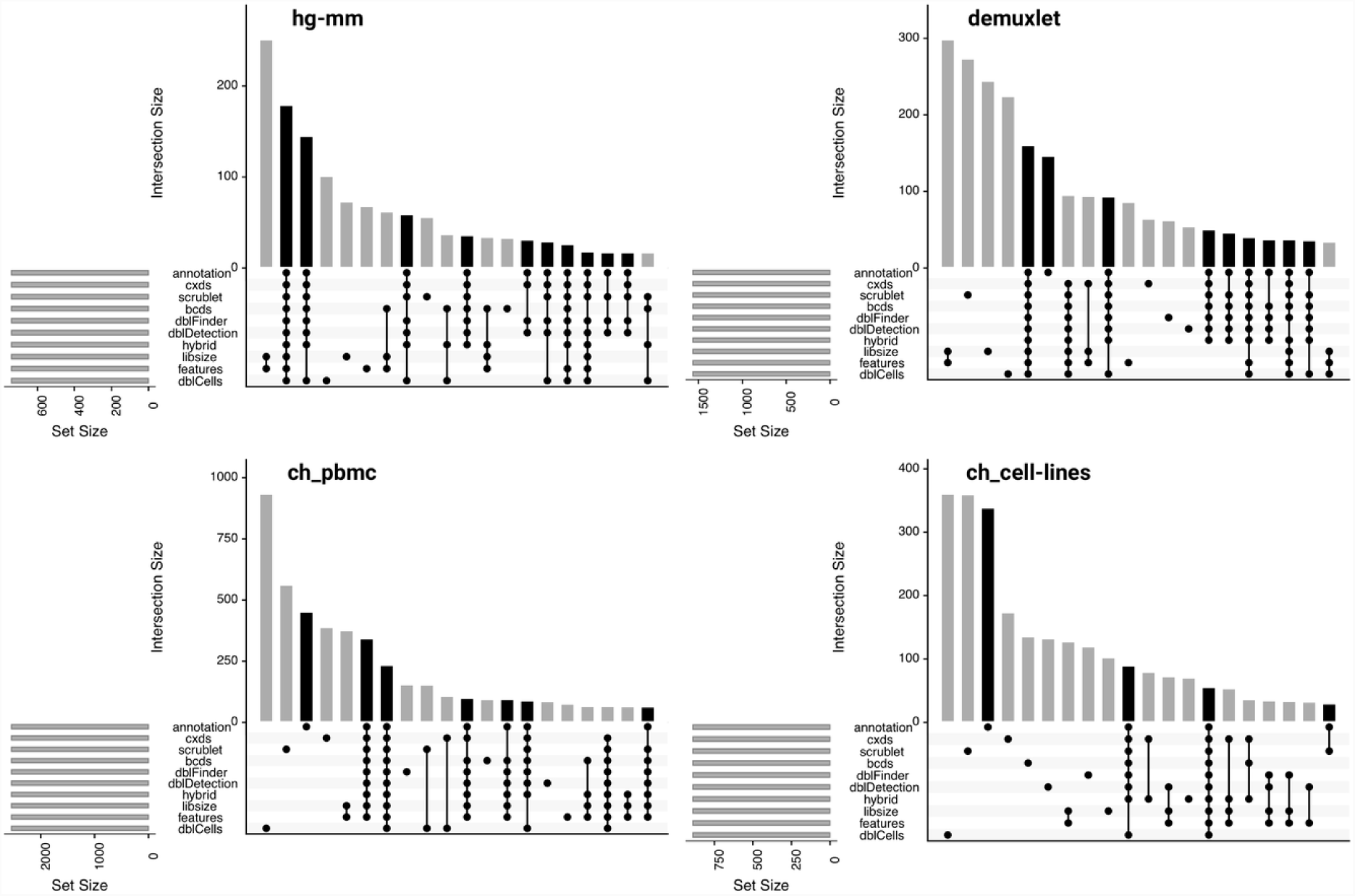
Comparison of doublet predictions. For four data sets (panels) we show upset plots (Conway *et al.*, 2017) comparing doublet predictions for nine prediction methods (including baseline methods) with annotated doublet cells. Bars showing the size of intersections containing annotated doublets (termed “annotation”) are in black, bars showing intersections without annotated doublets are in gray. We show the 20 largest intersection sets. For demuxlet, ch_pbmc and ch_cell-lines the set of doublets that gets missed by all prediction methods (i.e., consistent false negatives) is ranked number six, three and three in terms of size, respectively.

We find that in each data set (except `hg-mm`) there is a substantial number of annotated doublet cells that none of the computational annotation approaches recovers (black bars corresponding to the “annotation only” intersection). The `cxds`, `scrublet`, and `dblCells` methods often have a fairly large amount of FP predictions that are unique to the respective methods, as do `libsize` and/or `features`. While we note these differences, we also see that TP predictions are typically shared by many methods. In fact, with the exception of the `scrublet`-specific TP predictions in the `ch_cell-lines` data, all TP intersections have consistent predictions from at least four methods. That is, we observe better agreement between methods in terms of TP predictions as compared with FP predictions.

Further on, we compared the library size of cells, stratified by their annotations classes (TP, true negative (TN), FP, and false negative (FN) predictions) for each method and data set. Results are summarized in **Figure 4**. We see that for a few methods (`cxds, bcds, hybrid, dblDetection, dblFinder`) FP predictions tend to have higher library size compared with FN predictions, often comparable to TP predictions. Similar to what (McGinnis *et al.*, 2018) observe for `dblFinder`. We also find that this trend can vary for the same method between data sets (for example, `cxds` has this trend in all data sets except `ch_pbmc`, and `dblFinder` has it in all data sets except `hg-mm`).

**Figure 4:**
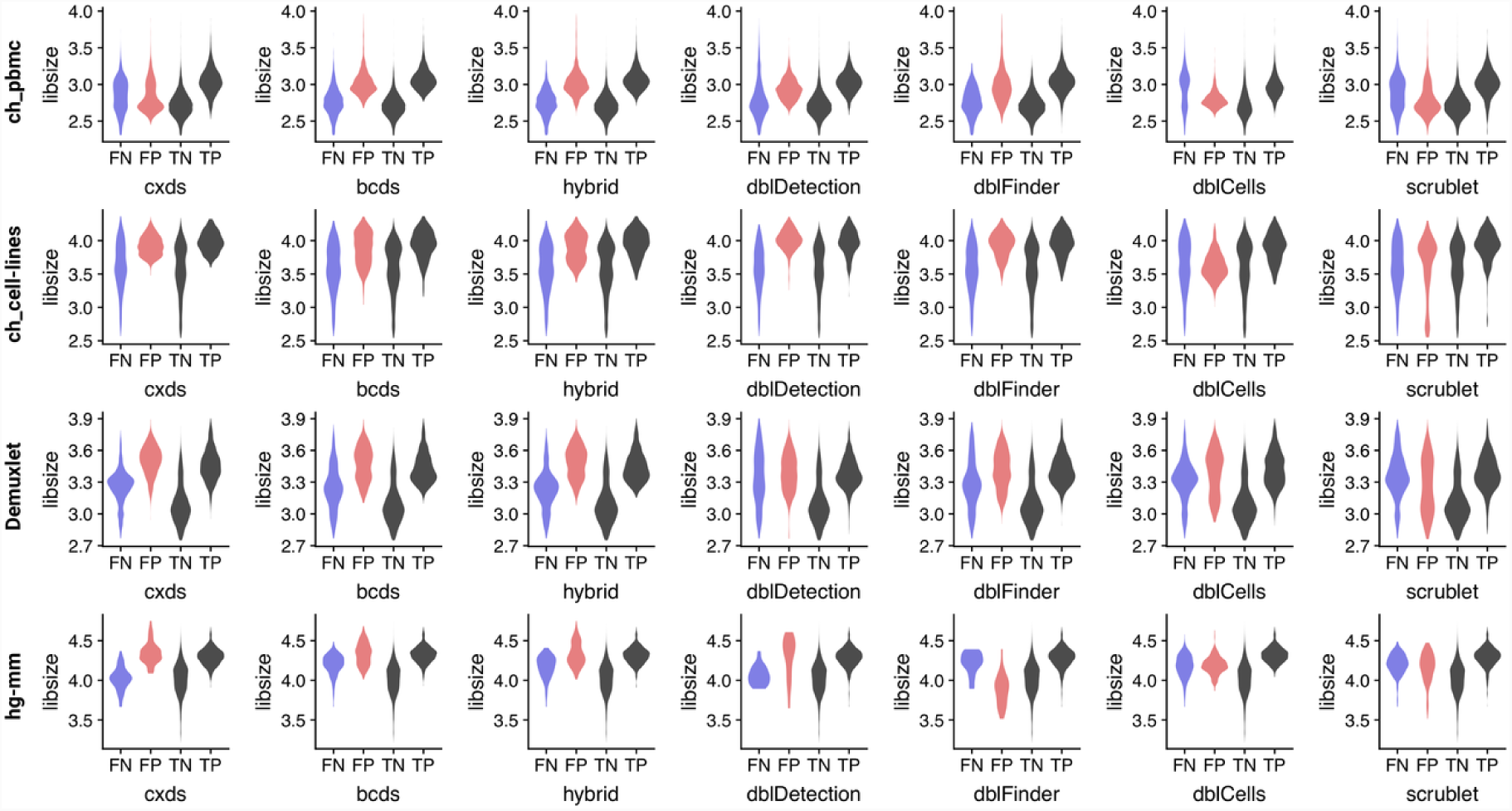
Library size of doublet annotations, stratified by prediction type. For each data set (rows) and each method (columns) violin plots of library sizes are shown for false negative predictions (FN, blue), false positive predictions (FP, red) and true negative and true positive predictions (TN and TP, both black). For most (but not all) method/dataset combinations library size in FP predictions tends to be higher compared with FN predictions.

Finally, the `ch_cell-lines` data set contains experimental annotation about whether a doublet is homotypic (from the same cell line) versus heterotypic. We used this to quantify the enrichment of TP predictions for heterotypic doublets across methods. Results are summarized in **Table 6**. We see that all methods (except `dblCells`) are significantly enriched for heterotypic doublets, with enrichment being most extreme for`cxds` and `bcds`. Next, we visually compared doublet annotations across methods and data sets.

#### Visual comparison of annotated doublets

We compared doublet predictions of each method for each data set in **Figure 5**. Rows correspond to computational annotation methods, and each major column represents a data set and is further sub-divided into four minor columns. The first minor column shows doublet scores with darker colors representing higher scores (i.e., more doublet-like barcodes). The second column shows TP predictions in the same coordinates, whereas the third and fourth columns show FP and FN predictions, respectively. The relative density for each type of prediction is indicated in color (TP: green, FP: red, FN: blue). As before, we choose cutoffs such that each method’s number of predicted doublets matches the number of experimentally annotated doublets.

**Figure 5:**
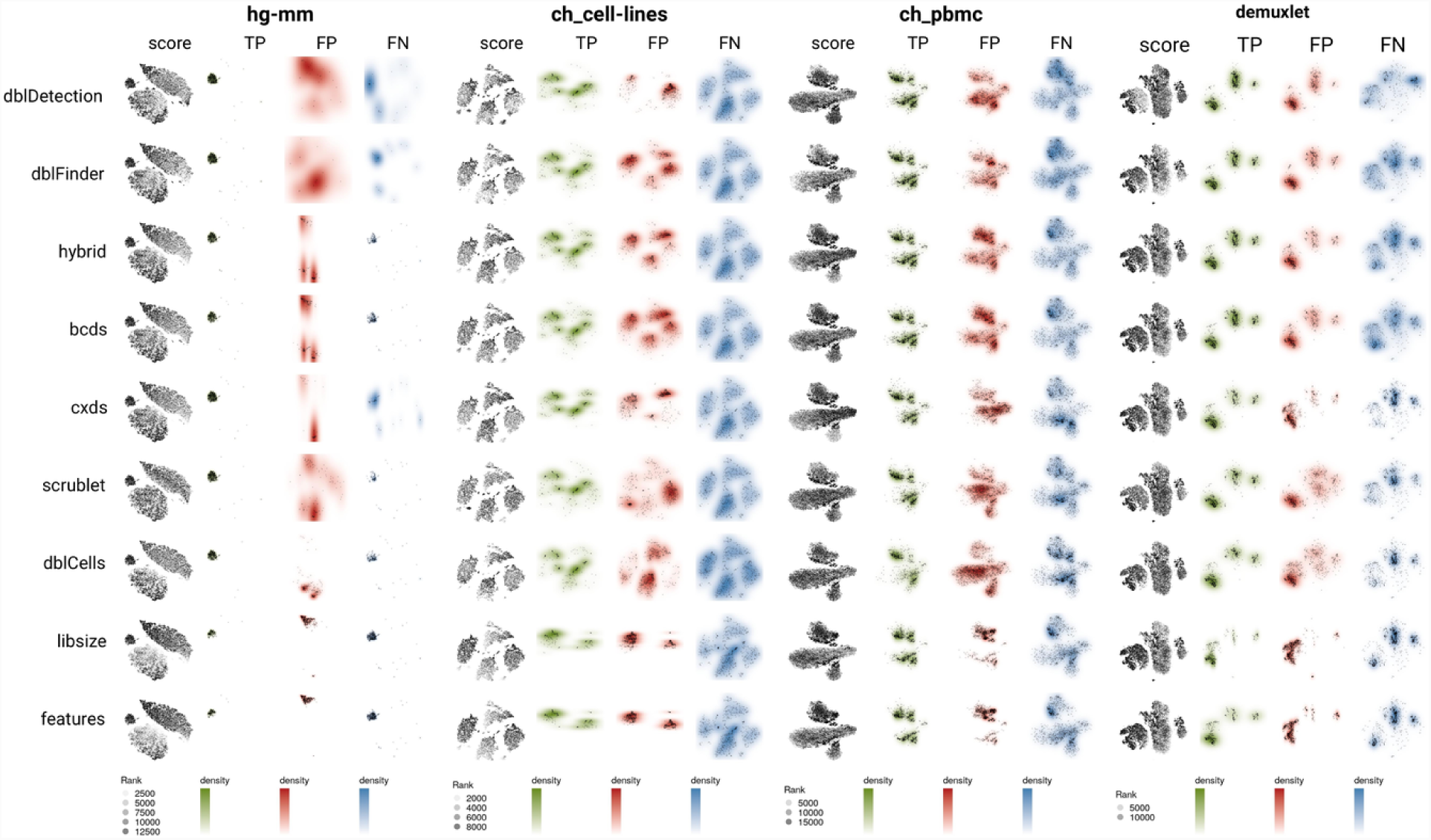
Visual comparison of doublet predictions. For each data set (major columns), we show four panels (minor columns) for the nine methods we compared (rows). The first left-most panel depicts all cells, shaded by the rank of the respective doublet prediction score for a method. The second, third and fourth panels show true positive (TP, green), false positive (FP, red) and false negative (FN, blue) predictions for a method, respectively. Shading in the respective color reflects the relative density, cells are shown in black.

For the `hg-mm` data, where computational annotations are mostly correct (Table 5), we see that TP and FN predictions are highly concentrated. However, we can still make out interesting differences between the methods in terms of where their FP predictions fall. `dblDetection`, `features` and `libsize` have FP predictions in similar areas for one type of cells pretty much exclusively, `scrublet` and `dblFinder` predict false positives more predominantly in the other cell type, while `hybrid` and `bcds` have FP predictions in both types of cells. For the `ch_cell-lines` data set, TP and FN predictions appear similar amongst non-baseline methods, while FP predictions appear distinct. For example, for `dblDetection` the FP density is highest in two of the four cell types (somewhat similar to the baseline methods), while for other methods FP predictions appear more broadly distributed. For the `ch_pbmc` data set we observe the biggest difference in terms of FP density between the two baseline methods and the rest, while for `demuxlet` we observe clearly visible differences in terms of TP, FP and FN density for all methods. For instance, FN predictions are more heterogeneous for the first four rows and more concentrated in the other methods. This coincides with higher FP concentration for the baseline methods and `cxds`, but not for `dblCells` and `scrublet`. Overall, we observe appreciable variability between doublet prediction methods, including the top three performers in **Table 2**, `dblFinder`, `hybrid` and `dblDetection`. This may suggest that none of the methods are close to optimal, and that an approach combining their respective strengths might further improve doublet annotation.

**Table 5:**
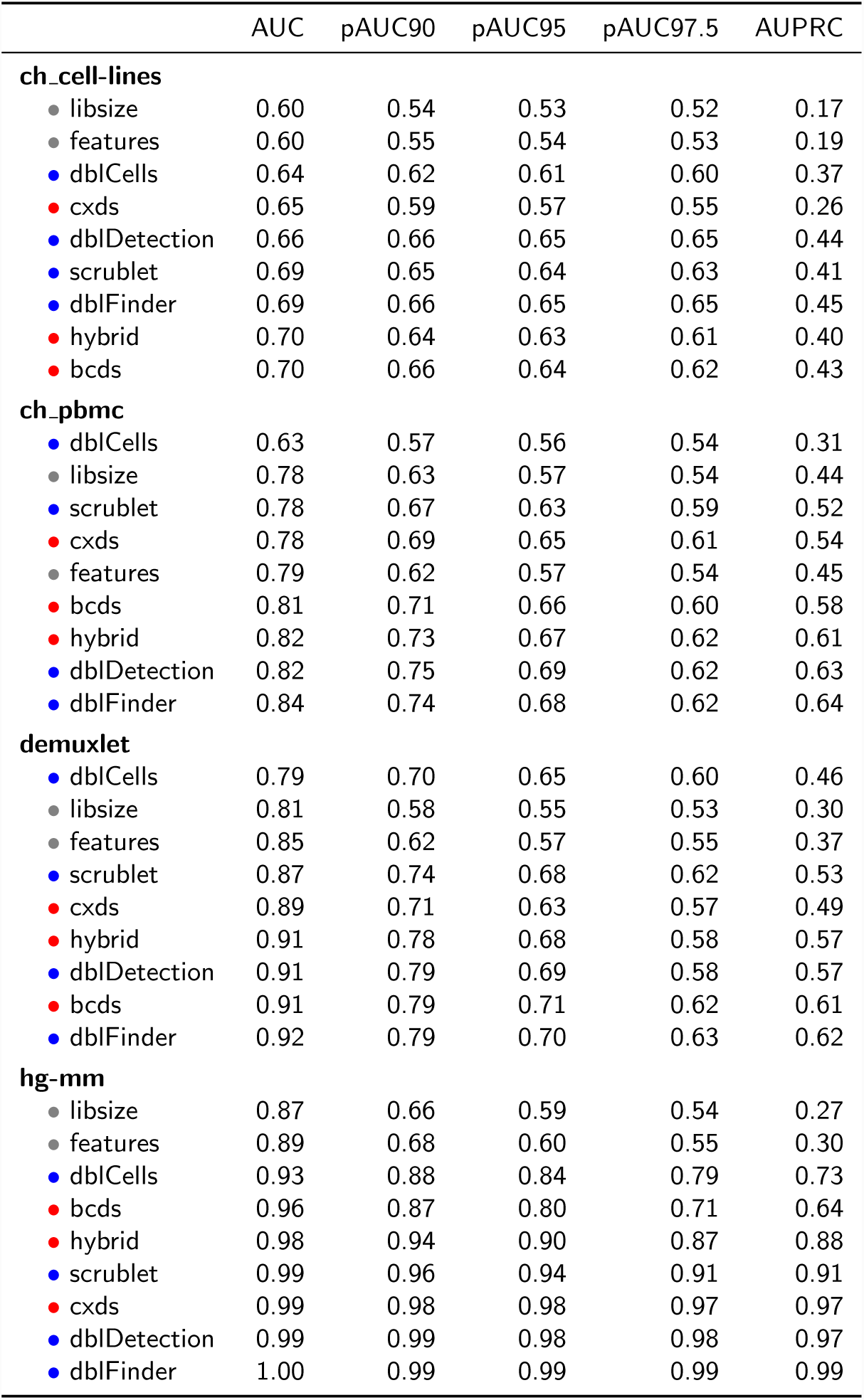
Performance of doublet prediction methods across four data sets. Gray bullets mark baseline methods, blue bullets mark current methods for doublet annotation, and red bullets mark proposed methods.

#### Running time comparison

We measured running times of the different methods we compare, and **Table 3** summarizes the results. We find that `cxds`, `bcds_7` (where we do not perform cross validation, see **Methods**) and `bcds` are able to annotate 10k cells in tens of seconds or on the order of a minute, while other methods take significantly longer. There is a distinct gap between “fast methods”, comprising the tools we propose and `scrublet`, and the rest. We note that computational doublet annotation can be performed for each chip/batch separately, and therefore we did not assess larger numbers of barcodes.

#### Comparison with dblDecon

Since`dblDecon` (DePasquale *et al.*, 2018) does not provide a doublet score, we could not include it in the previous analyses. To be able to still include it in our study, we applied it to all four data sets and and generated doublet predictions. For the `hg-mm` and `demuxlet` data sets, `dblDecon` did not annotate any doublets, and therefore we excluded them from this analysis. For the three remaining data sets and other tools in the comparison, we then generated the same number of annotated doublets as `dblDecon` by choosing appropriate score thresholds. **Table 8** summarizes results, with rows ordered by average precision across data sets. Surprisingly, we find that `dblDecon` does not perform well in this comparison, even though it determined the number of positive doublet calls for all methods. We also see a wide range of precision and sensitivity values across methods, while specificities are high due to the large amount of true negatives in all data sets.

## Discussion

We have introduced single cell doublet scoring, `scds`, encompassing three methods (`cxds`, `bcds`, and `hybrid`) for the *in silico* annotation of doublets in scRNA seq data. We have applied them to four data sets with experimental doublet annotations, and they all outperform baseline approaches. `cxds` is based on co-expression of gene pairs, and it is quite different from current approaches, because it does not utilize artificially generated doublets, and it works on a binarized absence/presence version of the RNA expression data. It features fast running times and provides users the opportunity to investigate pairs of genes driving doublet predictions (see **Methods** and **Results**). Binary classification based doublet scoring, `bcds`, is more similar to established methods and utilizes artificially generated doublets. However, in contrast to other tools (see **Introduction** for short descriptions), it does not make use of dimension reduction techniques, nor does it employ nearest neighbors for doublet scoring. Finally, `hybrid` is a combination of`cxds` and `bcds` that performs better than either method alone. In summary, our approaches are complementary to existing tools and work well for annotating doublets in scRNA-seq data.

We note that we do not estimate the number of doublets in a data set, but rather score cells/barcodes and rank them from most doublet-like to least doublet-like. Therefore, our annotations are most useful when an estimate about the expected doublet rate is available (for instance, 10X Genomics provides them in their *“User Guide for Chromium Single Cell 3’ Reagent Kits”*, based on the number of cells loaded on a chip), or when researchers wish to include a doublet score as only one of many factors in their decision about which cells may be excluded prior to downstream analyses. Our approaches share some conceptual limitations with other methods, which have been discussed in the literature (e.g, (Wolock *et* al., 2018)). Specifically, successful doublet identifications require that doublets are rare, that mixtures of more than two cells are even more rare, and that single cell instances of cell types in doublets are present in the data at appreciable frequency. Further on, our approaches are more sensitive towards identifying heterotypic doublets as compared with doublets comprised of two cells of the same type (also see **Table 6).**

**Table 6:** Enrichment of heterotypic doublets in true positive annotations. OR = odds ratio. P-values are via a Fisher exact test.

|  | cxds | bcds | hybrid | scrublet | dblFinder | dblDetection | libsize | numfeat | dblCells |
| --- | --- | --- | --- | --- | --- | --- | --- | --- | --- |
| OR | 7.70 | 5.39 | 3.97 | 3.32 | 3.19 | 3.04 | 2.62 | 1.20 | 0.98 |
| p-value | 7.38e-21 | 2.03e-16 | 2.29e-11 | 2.20e-10 | 6.82e-10 | 1.06e-07 | 2.46e-07 | 4.24e-01 | 9.18e-01 |

**Table 7:** Supplemental table. Comparison of annotation performance of `bcds_7` with `bcds` and `cxds`. We find that `bcds_7` performs comparable to `bcds`, but at a fraction of the running times (see Table 3).

|  | AUROC | pAUC90 | pAUC95 | pAUC97.5 | AUPRC |
| --- | --- | --- | --- | --- | --- |
| **ch_cell-lines** |  |  |  |  |  |
| bcds | 0.70 | 0.66 | 0.64 | 0.62 | 0.43 |
| cxds | 0.65 | 0.59 | 0.57 | 0.55 | 0.26 |
| bcds_7 | 0.69 | 0.65 | 0.63 | 0.61 | 0.41 |
| **ch_pbmc** |  |  |  |  |  |
| bcds | 0.81 | 0.71 | 0.66 | 0.60 | 0.58 |
| cxds | 0.78 | 0.69 | 0.65 | 0.61 | 0.54 |
| bcds_7 | 0.81 | 0.71 | 0.65 | 0.61 | 0.58 |
| **demuxlet** |  |  |  |  |  |
| bcds | 0.91 | 0.79 | 0.71 | 0.62 | 0.61 |
| cxds | 0.89 | 0.71 | 0.63 | 0.57 | 0.49 |
| bcds_7 | 0.90 | 0.77 | 0.69 | 0.61 | 0.58 |
| **hg-mm** |  |  |  |  |  |
| bcds | 0.96 | 0.87 | 0.80 | 0.71 | 0.64 |
| cxds | 0.99 | 0.98 | 0.98 | 0.97 | 0.97 |
| bcds_7 | 0.97 | 0.90 | 0.85 | 0.77 | 0.74 |

**Table 8:** Comparison of doublet annotation methods with dblDecon. TP: true positives, sen: sensitivity, spe: specificity, pre: precision

|  | ch_cell-lines |  |  |  | ch_pbmc |  |  |  |
| --- | --- | --- | --- | --- | --- | --- | --- | --- |
|  | TP | sen | spe | pre | TP | sen | spe | pre |
| dblDecon | 180 | 0.20 | 0.92 | 0.23 | 829 | 0.33 | 0.81 | 0.25 |
| dblCells | 280 | 0.31 | 0.93 | 0.36 | 915 | 0.36 | 0.82 | 0.28 |
| libsize | 163 | 0.18 | 0.92 | 0.21 | 1501 | 0.59 | 0.86 | 0.46 |
| features | 179 | 0.20 | 0.92 | 0.23 | 1513 | 0.59 | 0.86 | 0.46 |
| cxds | 240 | 0.27 | 0.93 | 0.31 | 1439 | 0.57 | 0.86 | 0.44 |
| scrublet | 312 | 0.35 | 0.94 | 0.40 | 1431 | 0.56 | 0.86 | 0.43 |
| bcds | 338 | 0.38 | 0.94 | 0.44 | 1680 | 0.66 | 0.88 | 0.51 |
| hybrid | 334 | 0.38 | 0.94 | 0.43 | 1712 | 0.67 | 0.88 | 0.52 |
| dblDetection | 328 | 0.37 | 0.94 | 0.42 | 1769 | 0.70 | 0.88 | 0.54 |
| dblFinder | 339 | 0.38 | 0.94 | 0.44 | 1806 | 0.71 | 0.89 | 0.55 |

We also compared our methods and four existing tools that provide doublet scores across four data sets, and we find appreciable heterogeneity between computational doublet annotation methods. No tool consistently outperforms all others, and performance varies between data sets. Our tools perform well, especially if running time is a consideration. Averaged across data sets, `dblDetection`, `hybrid` and `dblFinder` are the top performing methods (**Table 2**). Investigating doublet predictions of each method in more detail, we find that (i) for most data sets there is a sizable fraction of experimentally annotated doublets that is consistently missed by all methods, (ii) many correctly annotated doublets share the consensus of most methods, and (iii) methods differ mostly in terms of their false positive annotations, and these tend to be method-specific (i.e., typically not shared between methods). This implies that while methods differ in their doublet annotations (appreciable variability in terms of false positives), no method is yet able to recover a sizable fraction of annotated doublets (false negatives shared by all approaches). Therefore, we believe there is room to further improve computational doublet annotation. Specifically, we note that for `bcds` we used default parameters and did not really engage in parameter tuning, which could in principle lead to substantial improvements. The reason is that with only four doublet data sets available, we believe that there is some danger of inadvertent “information leak”, and therefore optimizing parameters may lead to overfitting. But this concern will decrease as more (and more diverse) data with experimental doublet annotations become available.

We note that results of our method comparisons necessarily depend on the data sets we used, and how they were processed. We attempted to minimize processing steps as much as possible, and we did not filter cells in addition to the original publications. We used published/provided experimental annotations, and for cell hashing (Stoeckius *et al.*, 2018) we followed the annotation strategy prescribed by the authors (see **Methods**). However, we are cognizant that alternative data processing strategies are equally reasonable and may have the potential to impact results. Further on, many analysis steps include random sampling in some way, thereby inducing a certain amount of stochasticity. Therefore we have made the code for our analyses available (https://github.com/kostkalab/scds_manuscript) and provide a docker container (https://hub.docker.com/r/kostkalab/scds) for other researchers. Finally, in our study we used experimentally annotated doublets as gold standard ignoring shortcomings of the respective experimental approaches, for example that some are not able to identify identically barcoded doublets (McGinnis *et al.*, 2018). However, in the absence of better experimental data we feel there are little alternatives to this approach.

In summary, *in silico* doublet annotation enriches single cell RNA sequencing data and can guard against over interpretation of results. From our comparison we find that current approaches (including ours) are able to annotate doublets more accurately than baseline methods, but also that there appears to be room for improvement as more data sets with experimental annotations become available. We introduced new light-weight methods for computational doublet annotation, which perform well in comparison to the status quo. They all feature comparably short running times, and co-expression based doublet scoring produces biologically interpretable results. Therefore, we provide researchers with new and useful tools to study and increase the value of their single cell RNA sequencing data.

## Acknowledgements

This research was supported by the National Institute of General Medical Sciences of the National Institutes of Health under award number R01GM115836, and by the University of Pittsburgh School of Medicine.

## Footnotes

1 We use the terms “cell” and “barcode” interchangeably, and sometimes use “doublet cells” to refer to barcodes coding for two or more cells.

## References

Li, H. et al. (2017) Reference component analysis of single-cell transcriptomes elucidates cellular heterogeneity in human colorectal tumors.. Nature genetics, 49, 708–718.

Segerstolpe, Å. et al. (2016) Single-Cell Transcriptome Profiling of Human Pancreatic Islets in Health and Type 2 Diabetes.. Cell metabolism, 24, 593–607.

Potter, S.S. (2018) Single-cell RNA sequencing for the study of development, physiology and disease.. Nature reviews. Nephrology, 14, 479–492.

Stegle, O. et al. (2015) Computational and analytical challenges in single-cell transcriptomics.. Nature reviews. Genetics, 16, 133–145.

Vallejos, C.A. et al. (2017) Normalizing single-cell RNA sequencing data: challenges and opportunities.. Nature methods, 14, 565–571.

AlJanahi, A.A. et al. (2018) An Introduction to the Analysis of Single-Cell RNA-Sequencing Data.. Molecular therapy. Methods & clinical development, 10, 189–196.

Kiselev, V.Y. et al. (2019) Challenges in unsupervised clustering of single-cell RNA-seq data.. Nature reviews. Genetics.

Proserpio, V. et al. (2016) Single-cell analysis of CD4+ T-cell differentiation reveals three major cell states and progressive acceleration of proliferation.. Genome biology, 17, 103.

Klein, A.M. et al. (2015) Droplet barcoding for single-cell transcriptomics applied to embryonic stem cells.. Cell, 161, 1187–1201.

Alles, J. et al. (2017) Cell fixation and preservation for droplet-based single-cell transcriptomics.. BMC biology, 15, 44.

Kang, H.M. et al. (2018) Multiplexed droplet single-cell RNA-sequencing using natural genetic variation.. Nature biotechnology, 36, 89–94.

Stoeckius, M. et al. (2018) Cell Hashing with barcoded antibodies enables multiplexing and doublet detection for single cell genomics.. Genome biology, 19, 224.

Gehring, J. et al. (2018) Highly Multiplexed Single-Cell RNA-seq for Defining Cell Population and Transcriptional Spaces.

McGinnis, C.S. et al. (2018) MULTI-seq: Scalable sample multiplexing for single-cell RNA sequencing using lipid-tagged indices.

Wolock, S.L. et al. (2018) Scrublet: computational identification of cell doublets in single-cell transcriptomic data.

Wang, Y.J. et al. (2016) Single-Cell Transcriptomics of the Human Endocrine Pancreas.. Diabetes, 65, 3028–3038.

Ibarra-Soria, X. et al. (2018) Defining murine organogenesis at single-cell resolution reveals a role for the leukotriene pathway in regulating blood progenitor formation.. Nature cell biology, 20, 127–134.

Rosenberg, A.B. et al. (2018) Single-cell profiling of the developing mouse brain and spinal cord with split-pool barcoding.. Science (New York, N.Y.), 360, 176–182.

Bach, K. et al. (2017) Differentiation dynamics of mammary epithelial cells revealed by single-cell RNA sequencing.. Nature communications, 8, 2128.

Ziegenhain, C. et al. (2017) Comparative Analysis of Single-Cell RNA Sequencing Methods.. Molecular cell, 65, 631–643.e4.

Krentz, N.A.J. et al. (2018) Single-Cell Transcriptome Profiling of Mouse and hESC-Derived Pancreatic Progenitors.. Stem cell reports, 11, 1551–1564.

Lun, A.T.L. et al. (2016) A step-by-step workflow for low-level analysis of single-cell RNA-seq data with Bioconductor. F1000Research, 5, 2122.

Shor, J. and Gayoso, A. (2019) DoubletDetection. GitHub repository.

McGinnis, C.S. et al. (2018) DoubletFinder: Doublet detection in single-cell RNA sequencing data using artificial nearest neighbors. bioRxiv, 352484.

DePasquale, E.A.K. et al. (2018) DoubletDecon: Cell-State Aware Removal of Single-Cell RNA-Seq Doublets.

Levine, J.H. et al. (2015) Data-Driven Phenotypic Dissection of AML Reveals Progenitor-like Cells that Correlate with Prognosis.. Cell, 162, 184–97.

Gong, T. and Szustakowski, J.D. (2013) DeconRNASeq: a statistical framework for deconvolution of heterogeneous tissue samples based on mRNA-Seq data.. Bioinformatics (Oxford, England), 29, 1083–1085.

R Core Team (2018) R: A Language and Environment for Statistical Computing R Foundation for Statistical Computing, Vienna, Austria.

Chen, T. and Guestrin, C. (2016) XGBoost: A Scalable Tree Boosting System. In, Proceedings of the 22nd ACM SIGKDD International Conference on Knowledge Discovery and Data Mining, KDD ’16. ACM, New York, NY, USA, pp. 785–794.

Chen, T. et al. (2019) xgboost: Extreme Gradient Boosting.

Hastie, T. et al. (2001) The Elements of Statistical Learning, Data Mining, Inference, and Prediction Corrected Printing, 2003. New York.

10X Genomics Human Mouse Cell Line Mixture.

Zerbino, D.R. et al. (2018) Ensembl 2018. Nucleic Acids Research, gkx1098–.

Durinck, S. et al. (2009) Mapping identifiers for the integration of genomic datasets with the R/Bioconductor package biomaRt. Nature Protocols, 4, 1184.

Butler, A. et al. (2018) Integrating single-cell transcriptomic data across different conditions, technologies, and species. Nature Biotechnology, 36, 411.

Allaire, J.J. et al. (2018) reticulate: Interface to ‘Python’.

McGinnis, C. (2018) DoubletFinder v2.0.0. GitHub repository.

Erichson, N.B. et al. (2016) Randomized Matrix Decompositions using R. arXiv preprint arXiv:1608.02148.

Krijthe, J.H. (2015) Rtsne: T-Distributed Stochastic Neighbor Embedding using Barnes-Hut Implementation.

Robin, X. et al. (2011) pROC: an open-source package for R and S+ to analyze and compare ROC curves. BMC Bioinformatics, 12, 77.

Keilwagen, J. et al. (2014) Area under Precision-Recall Curves for Weighted and Unweighted Data. PLoS ONE, 9, e92209.

Davis, J. and Goadrich, M. (2006) The relationship between Precision-Recall and ROC curves.

Conway, J.R. et al. (2017) UpSetR: an R package for the visualization of intersecting sets and their properties.. Bioinformatics (Oxford, England), 33, 2938–2940.

